# The dual role of the 16mer motif within the 3’ untranslated region of the variant surface glycoprotein of *Trypanosoma brucei*

**DOI:** 10.1101/2023.12.21.572740

**Authors:** Majeed Bakari-Soale, Christopher Batram, Henriette Zimmerman, Nicola G. Jones, Markus Engstler

## Abstract

The variant surface glycoprotein (VSG) of African trypanosomes is essential for survival of bloodstream form parasites. These parasites undergo antigenic variation, an immune evasion strategy in which they periodically switch VSG expression from one isoform to another. The molecular processes central to the expression and regulation of the VSG are however not fully understood. In general, the regulation of gene expression in trypanosomes is largely post-transcriptional. Regulatory sequences, mostly present in the 3’ UTRs, often serve as key elements in the modulation of the levels of individual mRNAs. In *T. brucei* VSG genes, a 16mer motif within the 3’ UTR has been shown to be essential for the stability of *VSG* transcripts and abundant VSG expression. This motif is 100 % conserved in the 3’ UTRs of all transcribed and non-transcribed VSG genes. As a stability-associated sequence element, the absence of nucleotide substitutions in the 16mer is however exceptional. We therefore hypothesised that the motif is involved in other essential roles/processes besides stability of the *VSG* transcripts.

In this study, we demonstrate that the 100 % conservation of the 16mer motif is not essential for cell viability or for the maintenance of functional VSG protein levels. We further show that the intact motif in the active VSG 3’ UTR is neither required to promote VSG silencing during switching nor is it needed during differentiation from bloodstream forms to procyclic forms. Ectopic overexpression of a second VSG, however, requires the intact 16mer motif within the ectopic VSG 3’ UTR to trigger silencing and exchange of the active VSG, suggesting a role for the motif in transcriptional VSG switching. The enigmatic 16mer motif therefore appears to play a dual role in transcriptional *VSG* switching and *VSG* transcript stability.

## Introduction

The African trypanosome *Trypanosoma brucei* is a unicellular protist parasite of medical and veterinary importance. The parasite is strictly extracellular and is transmitted between its mammalian hosts by an insect vector, the tsetse fly. In the mammalian host, *T. brucei* thrives in the blood and tissue fluids despite being exposed to the host’s immune system. Infection by the parasite can become chronic, lasting for months or years. The ability of the parasite to maintain chronic infections is due to constant changes in the identity of its surface coat through the process of antigenic variation (Vickerman, 1978; reviewed in Horn, 2014).

The *T. brucei* cell surface is covered by a dense layer of variant surface glycoprotein (VSG) composed of 10 million molecules linked to the cell surface via a glycophosphatidylinositol (GPI) anchor (Cross, 1975; Grünfelder *et al*., 2002; Schwede and Carrington, 2010). Each individual *T. brucei* cell expresses a single VSG out of a repertoire of over 2000 VSG isoforms (Cross, Kim and Wickstead, 2014). The actively expressed *VSG* occupies one of approximately 15 telomeric loci referred to as expression sites (ES) (Hertz-Fowler *et al*., 2008; Hutchinson, Glover and Horn, 2016). Only a single ES associated with an extranucleolar RNA polymerase I (RNA pol-I) transcription compartment, called the expression site body (ESB), is transcriptionally active in individual cells (Chaves *et al*., 1998; Navarro and Gull, 2001). The active VSG sub-compartment associates with a VSG exclusion (VEX) complex that mediates allelic exclusion, thus ensuring monoallelic VSG expression (Faria *et al*., 2019). The active VSG gene can however be exchanged with a silent VSG gene by either homologous recombination of a silent VSG into the active ES or transcriptional activation of a previously silent ES and silencing of the active ES (*in situ* switch) (Taylor and Rudenko, 2006; Alsford *et al*., 2012; McCulloch, Morrison and Hall, 2014). The mechanism of transcriptional VSG switching remains unclear.

VSG is the most abundant protein in the mammalian life cycle stages, comprising about 10 % of the total protein (Wang, Wang and Field, 2010). *VSG* mRNA is also the most abundant mRNA in blood stream trypanosomes with a half-life reported between 1 and 4.5 h (Ehlers, Czichos and Overath, 1987; Fadda *et al*., 2014; Ridewood *et al*., 2017). The high abundance of VSG protein and the very high stability of the *VSG* transcripts compared to other genes transcribed from the same ES has been attributed to RNA pol-I transcription and post-transcriptional control of VSG expression. *T. brucei* and related kinetoplastids have their genomes organised into long polycistronic transcription units. Most genes in these organisms are therefore constitutively transcribed into primary polycistronic transcripts, which are then processed into individual transcripts by a coupled *trans*-splicing and polyadenylation reaction (Lebowitz *et al*., 1993). The regulation of gene expression in trypanosomes is therefore largely post-transcriptional (Clayton, 2019). The untranslated regions (UTRs) of mRNAs play critical roles in the post-transcriptional regulation of gene expression. *Cis*-regulatory elements in the 3’ UTR in particular have been shown to be involved in the regulation of mRNA stability and translation efficiency in trypanosomes (Berberof *et al*., 1995; Hotz *et al*., 1997; Erben *et al*., 2014; Ridewood *et al*., 2017; Jojic *et al*., 2018). Two highly conserved motifs, an 8mer and 16mer, exclusively present in the 3’ UTRs of VSGs have been implicated in the post-transcriptional control of VSG expression. The 8mer motif is thought to be involved in a fail-safe mechanism that ensures destabilisation of accidently-produced VSG transcripts in procyclics (do Nascimento *et al*., 2021). The 16mer motif on the other hand has been shown to be essential for *VSG* mRNA stability (Berberof *et al*., 1995; Ridewood *et al*., 2017). Two mechanisms describing the involvement of the 16mer motif in maintenance of *VSG* mRNA stability have been proposed recently. Viegas *et al*. (2022) suggested that the 16mer is required for N6-methyladenosine (m^6^A) modification of the poly(A) tails of *VSG* mRNAs. This m^6^A modification prevents deadenylation and promotes *VSG* mRNA stability. do Nascimento *et al*. (2021) on the other hand identified and described the RNA binding protein CFB2 as a 16mer binding factor mediating *VSG* mRNA stability by recruitment of a stabilising translation-promoting complex.

The fact that the 16mer motif is 100 % conserved across all *T. brucei* strains and isolates has so far gone unnoticed. Therefore, we asked why a simple mRNA-stabilising motif of 16 nucleotides should not show a single polymorphism, even in silent, non-transcribed copies of VSG genes. This is highly unusual. There is a stability-associated 16 nucleotide sequence element in the 3’ UTR of procyclin transcripts, which encode the major insect stage cell surface protein of trypanosomes. This sequence motif is not as highly conserved (Hehl *et al*., 1994; Furger *et al*., 1997). Also, the mRNA motifs M23 and M24 in yeast and the miR-381 and miR-219 sequence motifs in humans have stability-associated functions but do not show 100 % conservation (Shalgi *et al*., 2005; Xie *et al*., 2005; Santa-Maria *et al*., 2015; Long *et al*., 2019).

Therefore, we hypothesise that there might be additional functions responsible for the 100 % preservation of the 16mer motif. This possibility was tested in an exclusion procedure. The 100 % conservation of the motif was not required for the expression of functional levels of VSG protein and hence is dispensable for cell viability. We also showed that the intact motif in the ES-resident VSG 3’ UTR is neither required for VSG silencing during switching nor is it needed during differentiation from bloodstream forms to procyclic forms. However, 100 % conservation of the 16mer motif within the 3’ UTR of an ectopic VSG was essential for triggering efficient VSG silencing and coat exchange during a transcriptional VSG switch. This additional role of the motif in the process of VSG *in situ* switching may be the driving force for its absolute conservation in *T. brucei* VSGs.

## Results

### Full conservation of the 16mer motif is not essential for expression of functional levels of VSG protein

An extensive mutation analysis of the *VSG121* 3’ UTR was performed. The mutants were stably integrated into the active expression site, upstream of the endogenous VSG221, to yield so-called double-expressor (DEX) cell lines (Figure S1A).

As anticipated, we found that the deletion of a segment from the 198bp-containing 3’ untranslated region (UTR) that includes either the complete 16mer motif (Δ39-198), or specific portions of the 16mer (Δ46-52 and Δ49-53), led to a significant reduction in VSG levels (see Figure S1A, B).

In contrast, inversion of the 8mer motif (Inv28-35) or mutation of the region between the 8mer and 16mer motifs (AC41-42TA) did not affect VSG expression levels.

However, we identified mutations of the 16mer motif (C61A and TGA46-48ACT), which did not significantly impact VSG production (Figure S1A - C). Thus, in principle, the 100 % conservation of the 16mer is not essential for parasite viability. This was confirmed by subsequent RNAi-mediated depletion of the endogenous VSG221 in DEX-parasites that featured a second VSG with a dysfunctional 16mer. Upon induction of RNAi these parasites revealed the typical lethal phenotype following pre-cytokinesis arrest that has been described for VSG-deficient trypanosomes. However, this detrimental phenotype was rescued if the 3’ UTR of the second VSG in the expression site contained one of the ‘tolerated’ mutations (Figure S2).

To confirm that parasites are viable with the tolerated mutation TGA46-48ACT (also referred as N46-48) in the 16mer motif, we generated single-expressor cells (Δ221^ES^121_N46-48_) harbouring this mutation from stable DEX cells (Figure 1A). Knock-out of the endogenous *VSG221* resulted in parasites expressing only VSG121 from the VSG221 expression site. As a control, we generated a cell line with the wild-type VSG121 3’ UTR (Δ221^ES^121_WT_). Both cell lines were viable. The parasites expressing the VSG with a mutated 16mer motif (Δ221^ES^121_N46-48_) exhibited a minor growth phenotype (Figure 1B). The Δ221^ES^121_WT_ cells had a population doubling time (PDT) of ∼ 9 h while the Δ221^ES^121_N46-48_ cells had a PDT of ∼ 11 h. The slowed growth was not a result of cell cycle defects (Figure 1C). *VSG* mRNA was however 43 % higher in the cells expressing a VSG with the wild-type 16mer motif (79 %) compared to the cells expressing a VSG with the mutated 16mer (36 %) though both cell lines had similar levels of VSG protein (Figure 1D, E). The tolerated mutation appears to affect *VSG* mRNA abundance, which may impact the rate of VSG protein production resulting in the observed growth phenotype. In the steady state, however, the VSG protein abundance was not significantly different between wild-type and mutant. On the one hand this shows that trypanosomes can live with less than 50 % of VSG mRNA. On the other hand, the experiment proves that the 100 % conservation of the 16mer motif is neither required for cell viability nor for production of functional levels of the VSG protein.

**Figure 1.**
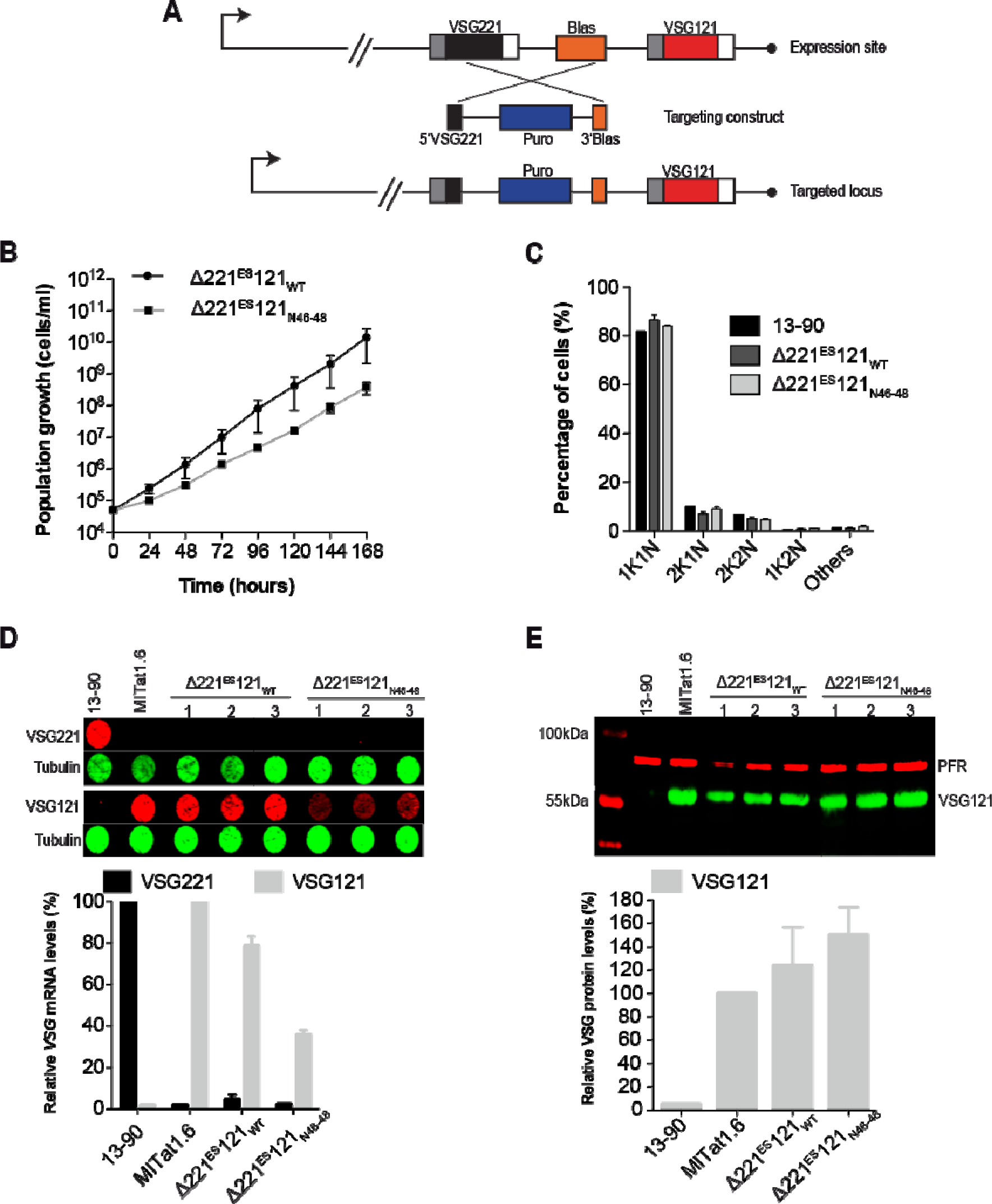
Substitution of the first three nucleotides of the 16mer motif supports expression of functional levels of VSG protein. (A) Schematic of the knockout strategy used in generating single-expressor cells from double-expressor cells with telomere proximal ectopic *VSG121*. The endogenous *VSG221* and blasticidin resistance (Blas) genes were replaced with a puromycin resistance gene (Puro). (B) Cumulative growth curve of single-expressor cells expressing *VSG121* with wild-type 16mer (Δ221^ES^121_WT_) and mutated 16mer (Δ221^ES^121_N46-48_). Data are averages (mean) from three independent clones with error bars representing the standard error of the mean (SEM). (C) Cell cycle analysis of Δ221^ES^121_WT_ and Δ221^ES^121_N46-48_ single-expressor cell lines. The number of nuclei and kinetoplasts were analysed microscopically in 200 DAPI-stained cells in each cell line. Three independent clonal cell lines were analysed, and data presented as mean percentages ± SEM. The parental 13-90 cell line was used as a control. (D) Quantification of *VSG* mRNA in Δ221^ES^121_WT_ and Δ221^ES^121_N46-48_ cells. *VSG221* and *VSG121* mRNA were quantified from RNA dot blots using fluorescently labelled probes normalised to *tubulin* mRNA (upper panel). *VSG221* and *VSG121* mRNA levels were expressed relative to levels in the parental 13-90 (VSG221) and MITat1.6 (VSG121) cells, respectively. Data are presented as mean ± standard error of the mean (SEM) of three independent clones. (E) Quantification of VSG protein in Δ221^ES^121_WT_ and Δ221^ES^121_N46-48_ cells. VSG121 protein levels were quantified from western blots and normalised to paraflagellar rod protein 1 and 2 (PFR) (upper panel). The levels of VSG121 protein were expressed relative to those found in the parental MITat1.6 cells. Data are presented as mean ± standard error of the mean (SEM) of three independent clones.

### An intact 16mer motif is not required for inclusion of N6-methyladenosine in poly(A) tails of *VSG* transcripts

It has been suggested that the 16mer motif is required for N6-methyladenosine (m^6^A) modification of *VSG* poly(A) tails contributing to stabilisation of the *VSG* transcripts (Viegas *et al*., 2022). To test whether this process requires the full 16mer motif, we performed m^6^A immunoblotting of poly(A)+ mRNA extracted from Δ221^ES^121_WT_ and Δ221^ES^121_N46-48_ cells. As a way of normalizing the m^6^A signal to initial transcript levels, we computed an ‘m^6^A index’ as described (Viegas *et al*., 2022) by dividing the relative intensity of the m^6^A signal in *VSG* transcripts by the corresponding *VSG* mRNA levels measured by quantitative dot blots (Figure 2). Amounts of *VSG121* mRNA were 20 % lower in the single-expressor cells expressing *VSG121* with a mutated 16mer (Δ221^ES^121_N46-48_) compared to cells expressing *VSG121* with the wild-type 16mer motif (Δ221^ES^121_WT_) (Figure 2A). The immunoblot revealed m^6^A bands corresponding to the *VSG121* transcript size in both Δ221^ES^121_WT_ and Δ221^ES^121_N46-48_ cells with m^6^A intensities of 0.43 and 0.33, respectively (Figure 2B, C). The m^6^A index was however identical (∼ 0.5 AU) in both Δ221^ES^121_WT_ and Δ221^ES^121_N46-48_ cells. This suggests that 100 % conservation of the 16mer motif is not required for m^6^A modification of *VSG* mRNA.

**Figure 2.**
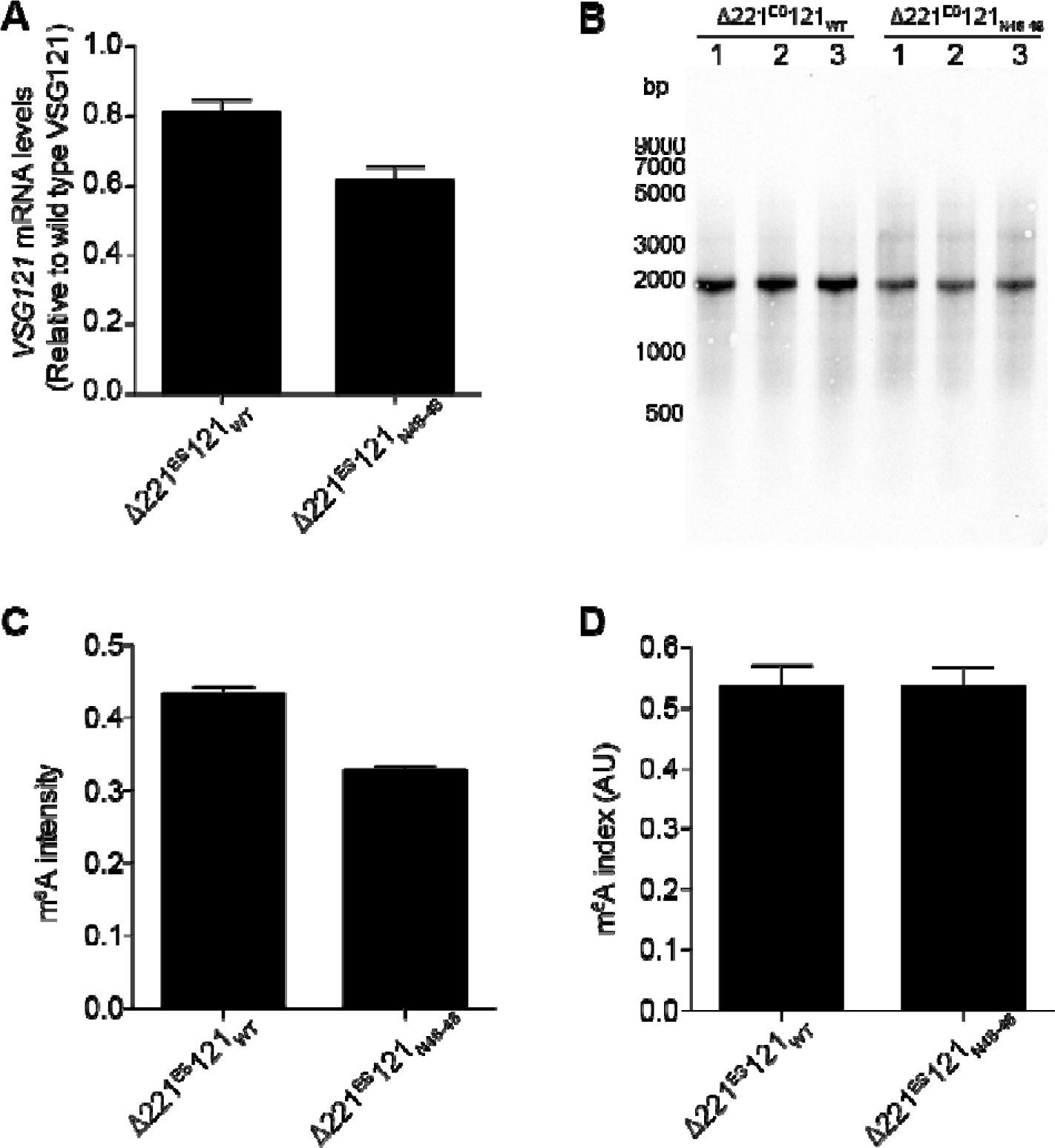
100 % conservation of the 16mer is not required for m^6^A modification of the poly(A) tails of *VSG* mRNA. (A) Quantification of *VSG* mRNA in Δ221^ES^121_WT_ and Δ221^ES^121_N46-48_ cells used in m^6^A immunoblots. *VSG121* mRNA was quantified from RNA dot blots, normalised to *tubulin* mRNA and expressed relative to the *VSG121* transcript levels in a MITat1.6 wild-type cell line. Data are presented as the mean ± standard error of the mean (SEM) of three independent clones each. (B) Immunoblots showing m^6^A amounts in mRNA from Δ221^ES^121_WT_ and Δ221^ES^121_N46-48_ cells. (C) Intensity of the m^6^A signal in *VSG121* mRNA measured using Image J. The intensity of the m^6^A in the *VSG121* band was expressed relative to the intensity of the entire lane. Data are presented as the mean ± standard error of the mean (SEM) of three independent clones each. (D) m^6^A index in the two single-expressor cell lines. The m^6^A index was computed as the ratio of the m^6^A intensity in (C) to the *VSG 121* mRNA levels in (A). Data are presented as the mean ± standard error of the mean (SEM) of three independent clones.

### An intact 16mer motif is not required for VSG silencing during differentiation from bloodstream form (BSF) to procyclic form (PCF)

Having established that the 100 % conservation of the 16mer is neither essential for expression of functional levels of VSG nor for the m^6^A modification of *VSG* mRNA, we next investigated a potential role of the conserved motif in VSG silencing during differentiation from BSF to PCF. This differentiation process is one of two events where destabilisation/silencing of the active ES occurs. The process can be activated *in vitro* by cold-shock and treatment with citrate and/or cis-aconitate (Brun and Schönenberger, 1981; Overath, Czichos and Haas, 1986; Engstler and Boshart, 2004). We performed the *in vitro* differentiation assay on Δ221^ES^121_WT_ and Δ221^ES^121_N46-48_ cells and monitored the *VSG121* mRNA and protein levels (Figure 3). Samples were collected 0, 6, 24 and 48 h after addition of 6 mM cis-aconitate and incubation of parasites at 27 °C. The *VSG121* mRNA and VSG121 protein levels were quantified using quantitative RNA and protein dot blots, respectively. A marked decrease of *VSG121* mRNA to less than 20 % of the wild-type levels was observed 6 h after induction of differentiation (Figure 3A). Less than 10 % of *VSG121* transcripts were present by 48 h post-induction. The decrease in mRNA was accompanied by a decrease in VSG protein to ∼ 25 % by 24 h (Figure 3B). The kinetics of silencing of the *VSG121* mRNA and VSG121 protein were similar in both cells expressing *VSG121* with either a wild-type or mutated 16mer motif, revealing that an intact 16mer motif is not essential for VSG silencing during differentiation. To directly investigate the effect of the 16mer-mutation on a possible crosstalk between the procyclin EP1-3’ UTR and the ES-resident VSG 3’ UTR, we integrated EP1-eYFP with a wild-type EP1 3’ UTR into a transcriptionally silent rDNA spacer in Δ221^ES^121_WT_ and Δ221^ES^121_N46-48_ cells to generate the cell lines Δ221^ES^121_WT_EP1^Tet^ and Δ221^ES^121_N46-48_EP1^Tet^, respectively (Figure S3). Tetracycline-induced overexpression of EP1-eYFP had no significant impact on the ES-resident VSG, irrespective of the *VSG121* harbouring either a mutated or the wild-type 16mer motif. This observation suggests that a potential crosstalk between the EP1 and VSG 3’ UTRs is not dependent on an intact 16mer motif.

**Figure 3.**
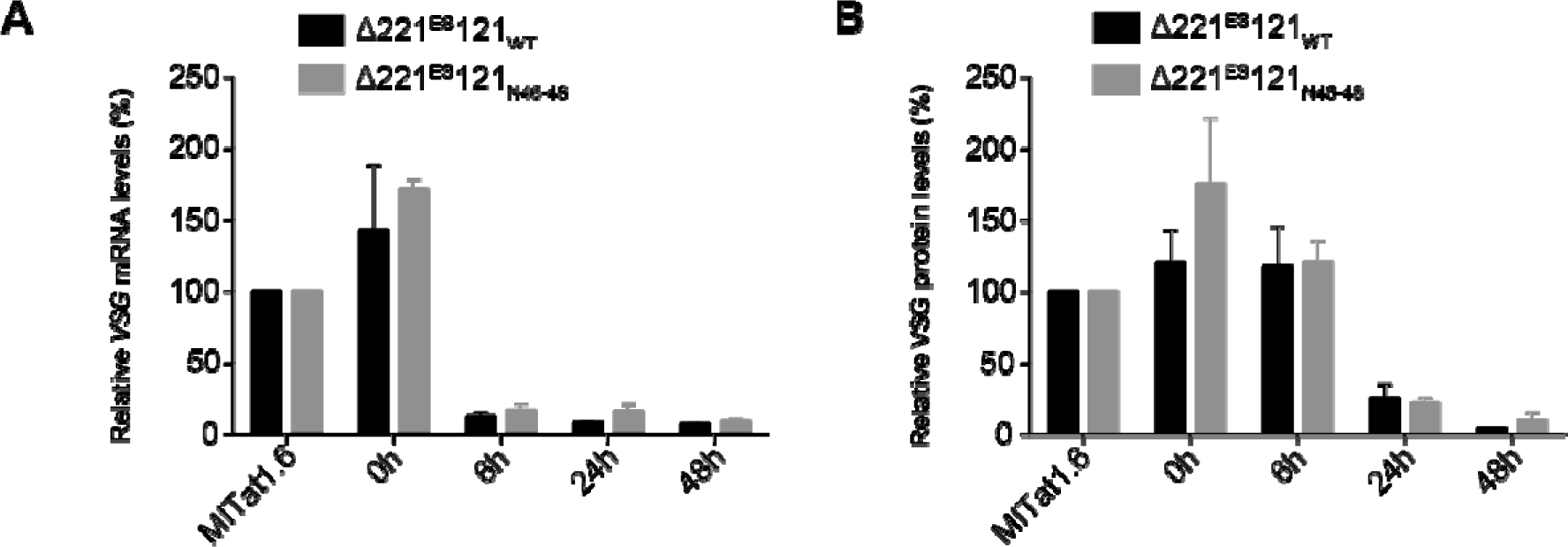
VSG silencing during differentiation from BSF to PCF occurs with similar kinetics in Δ221^ES^121_WT_ and Δ221^ES^121_N46-48_ cells. (A) *VSG121* mRNA levels in Δ221^ES^121_WT_ and Δ221^ES^121_N46-48_ cells during the course of differentiation. *VSG121* mRNA was quantified from RNA dot blots, normalised to *tubulin* mRNA and expressed relative to the *VSG121* transcript levels in a MITat1.6 wild-type cell line. Data are presented as the mean ± standard error of the mean (SEM) of three independent clones, respectively. (B) VSG121 protein levels in Δ221^ES^121_WT_ and Δ221^ES^121_N46-48_ cells during the course of differentiation. VSG121 protein was quantified from protein dot blots, normalised to PFR and expressed relative to the VSG121 protein levels in a MITat1.6 wild-type cell line. Data are presented as the mean ± standard error of the mean (SEM) of three independent clones, respectively.

### VSG silencing during *in situ* switching does not require an intact 16mer in the active VSG 3’ UTR

Apart from differentiation from BSF to PCF, VSG switching is the second event where silencing of the active VSG occurs in the parasite. Therefore, we next asked if the active VSG would have different silencing kinetics in the absence of the intact 16mer motif during switching. To test this possibility we used an inducible overexpression system that mimics transcriptional VSG switching (Batram *et al*., 2014) (Figure S4A). We integrated *VSG221* into a transcriptionally silent rDNA spacer in Δ221^ES^121_WT_ and Δ221^ES^121_N46-48_ cells. In the resulting Δ221^ES^121_WT_221^Tet^ and Δ221^ES^121_N46-48_221^Tet^ trypanosome lines, expression of the ectopic VSGs was driven by a tetracycline regulated T7 promoter. Upon induction of overexpression with tetracycline, the expression site resident *VSG121* transcript and protein levels decreased with similar kinetics in both the cell lines expressing *VSG121* with the wild-type or with the mutated 16mer motif, respectively (Figure 4). Within 24 h of overexpression, the ectopic VSG221 became the most abundant VSG expressed, similar to what we described in our earlier observations (Batram *et al*., 2014; Zimmermann *et al*., 2017). We also observed continuous growth in both cell lines after induction with Δ221^ES^121_WT_221^Tet^ cells having a PDT of ∼7.2 h compared to ∼8.4 h for uninduced cells, whereas the induced Δ221^ES^121_N46-48_221^Tet^ cells grew slightly faster with PDT of ∼7.1 h compared to ∼7.7 h for uninduced cells (Figure S4B).

**Figure 4.**
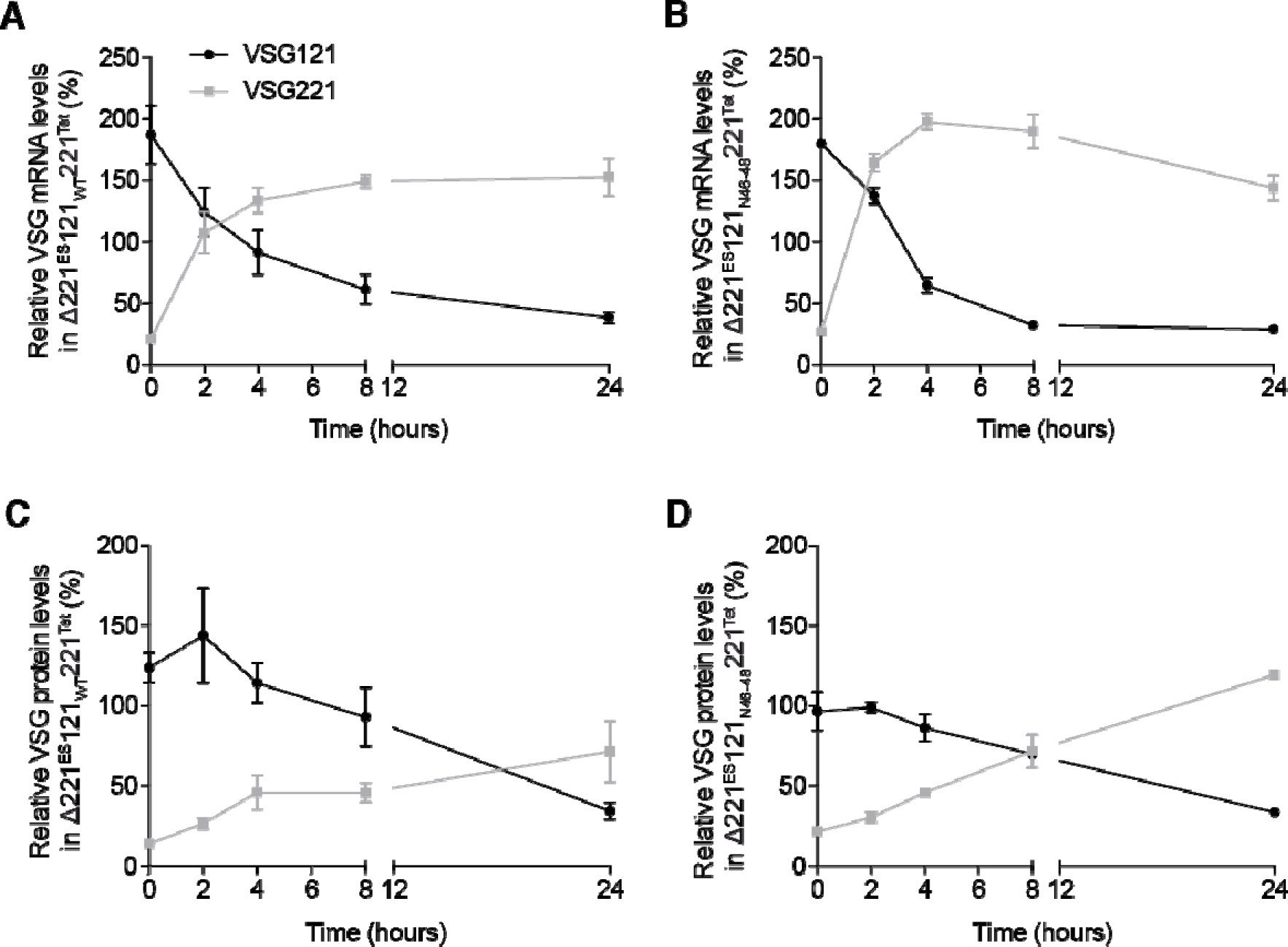
ES-resident VSG121 is silenced with similar kinetics in both the wild-type and mutated 16mer single-expressor cell lines upon ectopic *VSG221* overexpression. *VSG* mRNA (A, B) and VSG protein (C, D) monitored in Δ221^ES^121_WT_221^Tet^ and Δ221^ES^121_N46-48_221^Tet^ cells during the course of *VSG221* overexpression. *VSG* mRNA and VSG protein levels were quantified from RNA and protein dot blots, respectively. The *VSG* mRNA was normalised to *tubulin* mRNA and the protein amounts normalised to PFR. The VSG expression levels are given relative to levels in the parental 13-90 (VSG221) or MITat1.6 (VSG121) cells and expressed as mean ± standard error of the mean (SEM). Data presented are from three independent clones for each cell line.

### Ectopic overexpression of a second VSG requires an intact 16mer motif to trigger efficient VSG silencing and coat exchange

Since silencing of the VSG located at the expression site occurred independently of the presence of the conserved 16mer in its 3’ UTR, we wondered whether the 16mer motif was critical for expression of the "new" VSG. In our experiment, we mimicked this by inducible overexpression of a second VSG with either a complete 16mer or a mutant version. Would a 16mer 3’ UTR mutant efficiently induce VSG silencing and coat exchange as observed when VSG was overexpressed with wild-type 3’ UTRs? (Batram *et al*., 2014; Zimmermann *et al*., 2017). To test this, we integrated *VSG121* with the mutated 16mer 3’ UTR (VSG121_N46-48_) into a transcriptionally silent rDNA spacer in a parental cell line expressing VSG221. As a control, *VSG121* with an intact 16mer motif (VSG121_WT_) was used (Figure 5A). Overexpression of the ectopic wild-type VSG121_WT_ resulted in the expected severe growth arrest (Figure 5B) (Batram *et al*., 2014). Induction of the expression of the mutant VSG121_N46-48_, however, caused a much milder growth phenotype. This differential increase in population doubling time may be related to the degree of attenuation of the ES-resident VSG. Upon overexpression of wild-type VSG121_WT_, the endogenous *VSG221* transcripts are decreased by 65 % within 24 h, while induction of VSG121_N46-48_ resulted in slower kinetics and a less pronounced decrease of *VSG221* transcripts by about 40-45 % after 24 h. The difference observed cannot be attributed to the degree of VSG overexpression. With VSG121_N46-48_ overexpression, an increase of *VSG121* mRNA to ∼ 90 % of wild-type levels was observed within the first 2 h. There was a further increase to ∼ 120 % between 2 and 8 h, levelling to ∼ 90 % by 24 h. This was accompanied by a reduction in endogenous *VSG221* transcripts (Figure 5C). The endogenous *VSG221* transcripts however decreased to only 55 to 60 % within the first 24 h. This is in contrast to the kinetics of silencing observed upon overexpression of the wild-type VSG121_WT_, where the endogenous *VSG221* transcripts decreased to less than 35 % within the first 24 h (Batram *et al*., 2014). When the VSG121_N46-48_ mutant was overexpressed, the ectopic VSG121 protein reached only about 35 % of wild-type levels after 24 hours and decreased to less than 25 % by 72 hours (Figure 5D). The endogenous VSG221 protein remained abundantly expressed although higher amounts of *VSG121* transcripts were produced. Overexpression of the wild-type VSG121 on the other hand produced high levels of the ectopic VSG121 protein (∼ 74 % by 24 h) and caused a decrease of the endogenous VSG221 protein levels to ∼ 40 % by 24 h (Batram *et al*., 2014). These experiments indicate that overexpression of VSG121_N46-48_ was not effective in suppressing the expression of endogenous VSG221 at both the mRNA and protein levels. Thus, the last two experiments show that an intact 16mer is not required for the VSG gene to be silenced, however, the 16mer may be critical for the newly expressed VSG gene to silence the ES-resident VSG. These data therefore suggests that 100 % conservation of the 16mer is necessary to trigger efficient VSG silencing and coat exchange.

**Figure 5.**
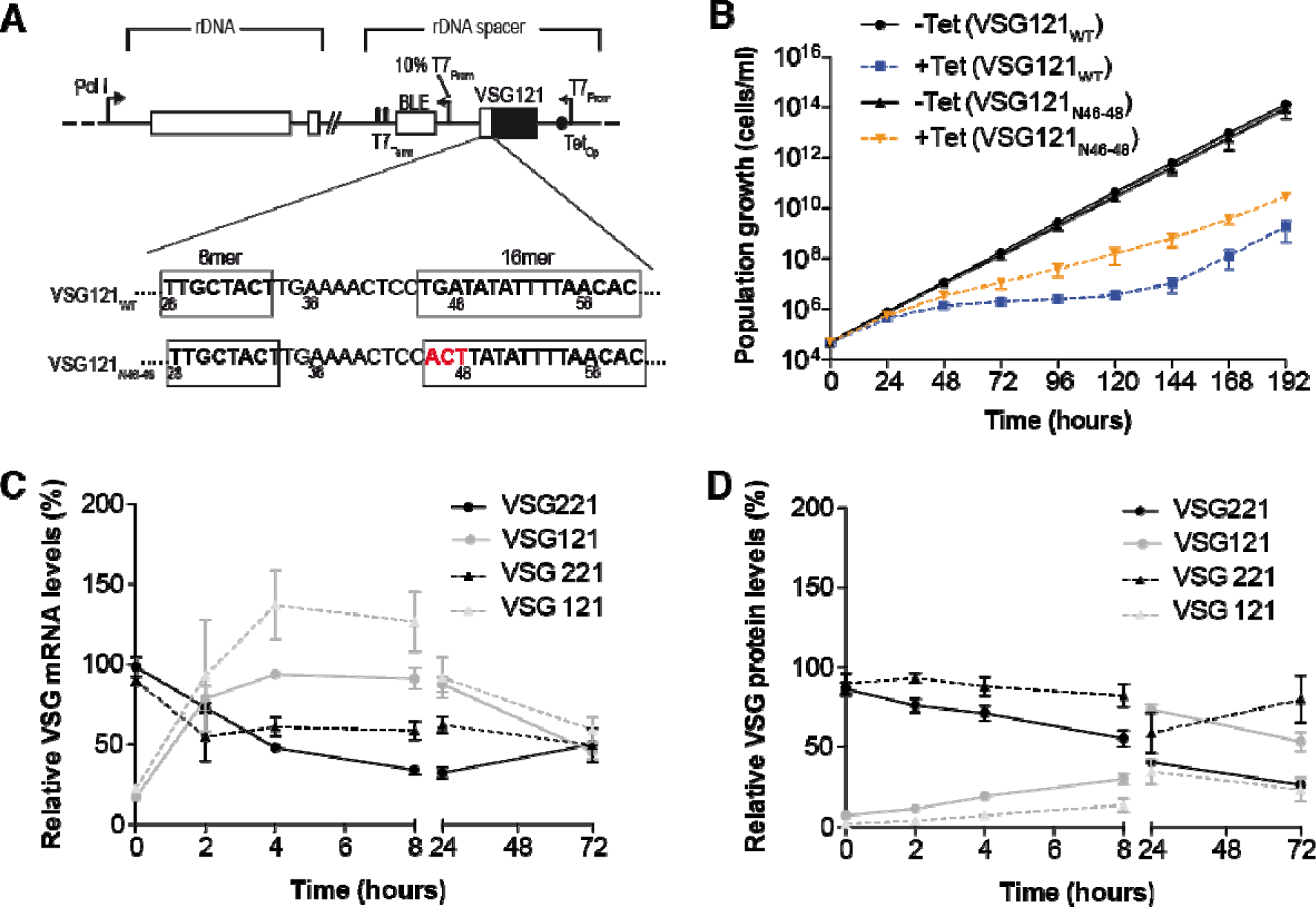
Overexpression of a second VSG with a mutated 16mer motif (VSG121_N46-48_) fails to trigger efficient silencing of endogenous VSG221. (A) Schematic showing the overexpression strategy of the ectopic *VSG121*. (B) Cumulative growth curves of cells before and after induction of VSG121_WT_ and VSG121_N46-48_ overexpression. Three independent clones were analysed for 8 days with or without 1 µg/ml tetracycline. *VSG* mRNA (C) and VSG protein (D) monitored upon induction of VSG121_WT_ and VSG121_N46-48_ overexpression. VSG levels from overexpression of VSG121_WT_ are indicated as circles connected by continuous lines whilst VSG levels from overexpression of VSG121_N46-48_ are indicated as triangles connected by dashed lines. The symbols and lines in black represent the endogenous VSG221 levels whilst the symbols and lines in grey represent the ectopic VSG121 levels. *VSG* mRNA and VSG protein levels were quantified from RNA and protein dot blots, respectively. The *VSG* mRNA was normalised to *tubulin* mRNA and the protein amounts normalised to PFR. The VSG expression levels are given relative to levels in the parental 13-90 (VSG221) or MITat1.6 (VSG121) cells and expressed as the mean ± standard error of the mean (SEM) of three independent clones, respectively.

### Less efficient VSG silencing is observed in slow growing cells upon overexpression of an ectopic VSG with a mutated 16mer motif

Our previous work had shown that overexpression of VSG121 causes VSG and ES attenuation, which is followed by dormancy of the parasite (Batram *et al*., 2014; Zimmermann *et al*., 2017). This phenotype was particularly striking in the monomorphic MITat 1.2 strain, while in the pleomorphic serodeme AnTat 1.1, *VSG* mRNA was similarly impaired, but ES attenuation and dormancy varied between mutant clones. Thus, we decided to challenge our observations with another VSG; we overexpressed VSG118 with either a wild-type (VSG118_WT_) or a mutated 16mer (VSG118_N46-48_) from the rDNA spacer. In fact, ectopic overexpression of VSG118_WT_ resulted in two phenotypically different types of clones. One set grew slightly faster than the uninduced cells (fast growers), while the other set grew slower (slow growers). VSG silencing and coat exchange occurred with similar kinetics in both slow and fast clones (Figure 6B-E). Overexpression of the mutant VSG118_N46-48_ also produced fast and slow growing clones (Figure 6A). In contrast to the overexpression of VSG118 with the wild-type 3’ UTR (VSG118_WT_), overexpression of the mutant VSG118_N46-48_ caused a much less efficient VSG silencing specifically in the slow growing clones (Figure 6B - E). This result supports our finding that an ectopic VSG requires an intact 16mer motif for efficient silencing of the ES-resident, endogenous VSG.

**Figure 6.**
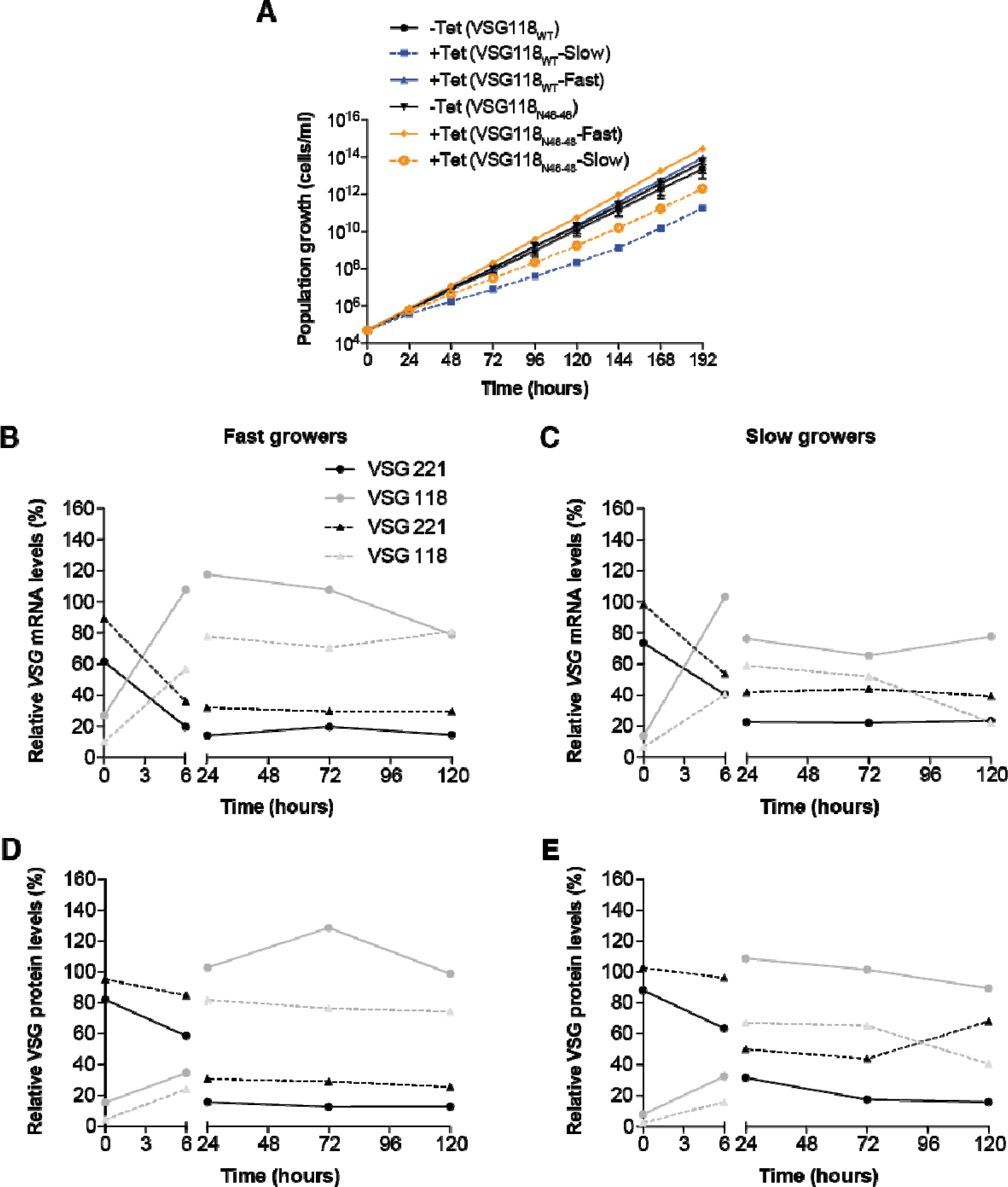
Endogenous VSG silencing is less efficient in slow growing cells obtained from VSG118 overexpression. (A) Cumulative growth curves after induction of ectopic VSG118_WT_ and VSG118_N46-48_ overexpression. Two distinct clonal populations (fast growers and slow growers) were observed. The plot for uninduced (-Tet) cells is from four independent clones and plots for the induced cells (+Tet) are from two clones each. Values are presented as mean. *VSG* mRNA levels in fast growing clones (B) and slow growing clones (C) during ectopic overexpression of VSG118_WT_ and VSG118_N46-48_. VSG protein levels in fast growing clones (D) and slow growing clones (E) during ectopic overexpression of VSG118_WT_ and VSG118_N46-48_. VSG levels from overexpression of VSG118_WT_ are indicated as circles connected by continuous lines whiles VSG levels from overexpression of VSG118_N46-48_ are indicated as triangles connected by dashed lines. The symbols and lines in black represent the endogenous VSG221 levels whiles the symbols and lines in grey represent the ectopic VSG118 levels. *VSG* mRNA and VSG protein levels were quantified from RNA and protein dot blots, respectively. The *VSG* mRNA was normalised to *tubulin* mRNA and the protein amounts normalised to PFR. Values were expressed relative to levels in the parental 13-90 (VSG221) or MITat1.5 wild-type cells (VSG118) and are presented as mean.

## Discussion

In this study we demonstrate that the 100 % conserved 16mer motif within the VSG 3’ UTR of *T. brucei* plays a role in VSG switching by facilitating efficient silencing and exchange of the active VSG. The VSG 3’ UTR has been previously shown to be important for stabilisation of *VSG* mRNA using a CAT reporter gene and VSG double-expressor cells (Berberof *et al*., 1995). Mutational analysis identified the 100 % conserved 16mer motif as the sequence element responsible for maintenance of the mRNA stability. Scrambling of the 16mer motif caused a massive reduction in the *VSG* mRNA half-life from over 100 min to ∼ 20 min (Ridewood *et al*., 2017). Prior to our study, the high conservation of the motif has been attributed solely to its role in transcript stability. As a stability-associated motif, it is however unusual that no nucleotide substitutions are present in this motif as seen in other stability-associated motifs. Using mutational analysis in stable double-expressor cells we confirmed that the 16mer motif is indeed essential for *VSG* mRNA stability. We however found a tolerated mutation of the motif involving substitution of up to three nucleotides (TGA to ACT). Introduction of this mutation into the 16mer motif of transgenic *T. brucei* cell lines expressing a single VSG resulted in production of VSG amounts sufficient for cell viability. These single-expressor cells expressing *VSG121* with a mutated 16mer transcribed 20 % to 43 % less *VSG121* mRNA compared to control cells (single-expressor cells expressing *VSG121* with wild-type 16mer). The cells expressing *VSG121* with a mutated 16mer motif also grew slower compared to the single-expressor cells expressing *VSG121* with the wild-type 16mer. VSG121 protein levels were however similar and comparable to wild-type amounts in both cell lines, suggesting that the intact 16mer motif is necessary for mRNA stability but not required for expression of functional levels of VSG protein. As the rate of synthesis of a protein is dependent on the concentration and translation efficiency of its mRNA (Polymenis and Aramayo, 2015), the cells expressing *VSG121* with the 16mer mutation may take longer to translate functional levels of the VSG protein due to the lower mRNA levels, hence the slower growth phenotype. Our data therefore showed that the 100 % conservation of the 16mer motif is not essential for functional VSG protein production, hence the high conservation of the motif may not solely be due to its role in transcript stability.

Two mechanisms have been proposed for the maintenance of the stability and abundance of the *VSG* transcripts. One of these mechanisms proposes the presence of a 16mer binding factor which prevents degradation of the mRNA by blocking association of the transcript with nucleases (Ridewood *et al*., 2017). CFB2 has been identified as the 16mer binding factor and further shown to act by recruiting a stabilising complex that includes MKT1, PBP1, PABP2, and the cap-binding translation initiation complex EIF4E6/G5 (do Nascimento *et al*., 2021; Bravo Ruiz *et al*., 2022). The second proposed mechanism involves a 16mer dependent m^6^A modification of poly(A) tails of VSG transcripts which inhibits RNA degradation by blocking deadenylation (Viegas *et al*., 2022). We investigated whether reduced m^6^A modification due to absence of the intact 16mer motif was responsible for the lower VSG transcript levels in the single-expressor cell lines expressing *VSG121* with a mutated 16mer. Analysis of the m^6^A modification in *VSG121* mRNA from the cells expressing *VSG121* with a mutated 16mer and cells expressing *VSG121* with wild-type 16mer revealed similar m^6^A levels, suggesting that 100 % conservation of the 16mer motif is not required for m^6^A modification of *VSG* transcripts. This does not however rule out the role this modification plays in transcript stability as subtle differences in m^6^A levels may not be detectable with the northwestern blot approach.

During a VSG switching event or differentiation of parasites from bloodstream forms to procyclic forms, the active VSG gene is silenced. This occurs due to repression of the active VSG with the expression of a new VSG or procyclin (Myler *et al*., 1984; Gruszynski *et al*., 2006). Destabilisation of the active *VSG* mRNA thus occurs. In the case of differentiating cells, the half-life of the *VSG* mRNA is reduced from ∼ 4.5 h in bloodstream trypanosomes to 1.2 h in the transforming cells (Ehlers, Czichos and Overath, 1987). Since the 16mer motif is an essential stability-associated motif, the role of its 100 % conservation in VSG silencing during VSG switching and differentiation from BSF to PCF were investigated. We found that regardless of the presence or absence of an intact 16mer motif in the active VSG 3’ UTR, silencing of the active VSG occurs rapidly and efficiently during switching or differentiation. This further suggests that 100 % conservation of the 16mer motif within the active VSG 3’ UTR is not essential for silencing of the active VSG during parasite differentiation and during a transcriptional VSG switch.

A transcriptional/*in situ* VSG switch involves activation of a new ES and silencing of the previously active ES. Therefore, having established that the 100 % conservation of the 16mer motif within the active VSG is not essential for silencing, we investigated whether the intact 16mer is required in the new VSG to trigger efficient silencing and exchange of the active VSG during switching. Using the overexpression system mimicking transcriptional VSG switching (Batram *et al*., 2014), we found that in the presence of an intact 16mer motif in the ectopic VSG121, the endogenous VSG221 was efficiently silenced and replaced with the ectopic VSG121 within 24 h of overexpression. This agreed with previous studies which have established that high level expression of an ectopic wild-type VSG triggers efficient silencing and exchange of the active VSG (Batram *et al*., 2014; Zimmermann *et al*., 2017). In the absence of the 100 % conserved 16mer motif however, the endogenous *VSG221* mRNA only decreased by 40 – 45 % within the first 24 h and this VSG remained the most abundantly expressed protein. Mutation of the 16mer motif therefore prevented efficient silencing and exchange of the endogenous VSG, suggesting that the 100 % conservation of the 16mer motif is necessary to trigger efficient VSG silencing and coat exchange during a transcriptional VSG switch. This finding was confirmed with the overexpression of VSG118_WT_ (with an intact 16mer motif) and VSG118_N46-48_ (with a mutated 16mer motif). Overexpression of the VSG118_WT_ produced two distinct growth phenotypes but regardless of the growth phenotype, the endogenous VSG221 was efficiently silenced and exchanged. Overexpression of VSG118_N46-48_ however produced contrasting results between the two clonal populations. In the fast growers, the endogenous VSG221 was efficiently silenced at both the mRNA and protein levels. In the slow growers however, the endogenous VSG221 was not efficiently silenced. The *VSG221* mRNA levels decreased to about 42 % by 24 h and the protein levels remained above 50 %. This data therefore confirms that the 100 % conservation of the 16mer motif is important for VSG silencing and coat exchange. The high conservation of the motif appears to be essential in a silent VSG for transcriptional switching between the active VSG and a previously silent VSG. This finding supports the assumption that the mechanism of antigenic switching operates differently in *T. brucei* compared to *T. congolense* and *T. vivax.* Although these other African trypanosome species possess the VSG, they lack the conserved 16mer motif and may therefore have evolved different sequence elements for regulation of stability and transcriptional VSG switching.

Our study has led to the identification of a previously undescribed role of the intact 16mer motif, as the 100 % conservation of the motif was found to be essential for triggering efficient VSG silencing and coat exchange during a mimicked *in situ* VSG switch. This additional role of the 16mer motif in antigenic variation, which is an important process in the parasite, may be the driving force for the 100 % conservation of the motif in all *T. brucei* VSG isoforms.

## Experimental procedures

### Trypanosome cultivation

All the cell lines generated in this study are based on monomorphic *T. brucei* Lister 427 13-90 cells (Wirtz *et al*., 1999). Monomorphic *T. brucei* Lister 427 MITat1.6 and *T. brucei* Lister 427 MITat1.5 wild-type cells were cultivated for use as controls in experiments. The cells were cultivated in HMI-9 medium (Hirumi and Hirumi, 1989) supplemented with 10 % heat-inactivated fetal calf serum (FCS) (Sigma-Aldrich, St. Louis, USA), at 37 °C and 5 % CO_2_. Previously transfected plasmids in *T. brucei* 13-90 cells were maintained by the addition of 5 µg/ml hygromycin and 2.5 µg/ml G418. Cell numbers were monitored by counting with a haemocytometer or Z2 Coulter counter (Beckman Coulter). For transfections to generate transgenic cell lines, 3 x 10^7^ *T. brucei* 13-90 cells were electroporated with 10 µg of linearised plasmid DNA using the Amaxa Basic Parasite Nucleofector Kit 1 and Nucleofector II device (program X-001) (Lonza, Switzerland).

### Plasmids and generation of transgenic trypanosome cell lines

For the generation of stable double-expressor (DEX) cell lines where the ectopic *VSG121* was integrated upstream of the endogenous *VSG221*, the *VSG121* gene with either a wild-type *VSG121* 3’ UTR or mutated *VSG121* 3’ UTRs was cloned into the plasmid pKD4f and transfected into *T. brucei* 13-90 cells. The intermediate plasmids pBSK.M1.6_WT_ and pBSK.M1.6_Δ116-198_ were generated by PCR amplification of *VSG121* from genomic DNA of *T. brucei* MITat1.6 wild-type cells. The PCR products were digested with NcoI and BamHI and ligated into the plasmid p2084 (MITat1.6T434A in pKD4 vector) digested with the same enzymes to generate the resultant plasmids. For the plasmids pBSK.M1.6_Δ63-198_, pBSK.M1.6_Δ39-198_ and pBSK.M1.6_Δ1-198_, the plasmid pBSK.M1.6_WT_ was used as template.

The PCR products were digested with NcoI and BamHI and ligated into the plasmid p2084 digested with the same enzymes to generate the resultant plasmids. The *VSG121* gene with either a wild-type or mutated 3’ UTR was subcloned into the pKD4f plasmid after EcoRI digest to yield pES-M1.6_WT_, pES-M1.6_Δ116-198_, pES-M1.6_Δ63-198_, pES-M1.6_Δ39-198_ and pES-M1.6_Δ1-198_. pBSK.M1.6_Δ46-52_ and pBSK.M1.6_Δ49-53_ were amplified in two separate PCR steps using the plasmid p2084 as template. The resulting PCR products were digested with EcoRI and BamHI and ligated into the pBSK plasmid backbone. The PCR products were then excised from the resulting plasmids with SpeI and HincII and subcloned into the pKD4f plasmid to generate pES-M1.6_Δ46-52_ and pES-M1.6_Δ49-53_. For plasmids pBSK.M1.6_Inv28-35_, pBSK.M1.6_AC41-42TA_, pBSK.M1.6_C61A_ and pBSK.M1.6_TGA46-48ACT_, a fusion PCR approach as described by (Heckman and Pease, 2007) was used to generate the VSG121 3’ UTR mutations. The resulting PCR products were ligated into pBSK plasmid backbones after digestion with EcoRI and BamHI. The *VSG121* CDS with mutated 3’ UTRs were subcloned into pKD4f plasmid after EcoRI digest to yield pES-M1.6_Inv28-35_, pES-M1.6_AC41-42TA_, pES-M1.6_C61A_ and pES-M1.6_TGA46-48ACT_. The pES plasmids were linearised with AvrII and KpnI and transfected into *T. brucei* 13-90 cells to produce the VSG121 double-expressor cell lines (WT, Δ116-198, Δ63-198, Δ39-198, Δ1-198, Δ46-52, Δ49-53, Inv28-35, AC41-42TA, C61A and TGA46-48ACT).

To generate the VSG121 single-expressor cell lines Δ221^ES^121_WT_ and Δ221^ES^121_N46-48_, double-expressor cell lines (221^ES^121_WT_ and 221^ES^121_N46-48_), where the ectopic *VSG121* was integrated downstream of the endogenous *VSG221*, were first generated. The endogenous *VSG221* was then knocked out using the plasmid pJET1.2_M1.2-Blas_KO. The double-expressor cell lines 221^ES^121_WT_ and 221^ES^121_N46-48_ were generated by transfecting the plasmids pbRn6.M1.6_WT_ and pbRn6M1.6_TGA46-48ACT.nPPT_, both linearised with SacI and SalI into *T. brucei* 13-90 cells, respectively. The plasmid pbRn6.M1.6_WT_ was generated as described by (Aroko *et al*., 2021). pbRn6M1.6_TGA46-48ACT.nPPT_ on the other hand was generated by mutagenesis PCR using the fusion PCR approach. Briefly, primer pairs C14/HZ35 and C15/HZ32 were used to amplify two PCR fragments from the template pbRn6M1.6_198. The two fragments were joined together in an additional PCR step using the primer pair HZ35/HZ32. The resultant PCR product was cloned into pJET1.2 to yield pJET1.2M1.6_TGA46-48ACT_. The M1.6_TGA46-48ACT_ fragment was then excised with HindIII and MVa1269I and ligated into the pbRn6 backbone obtained from digestion of pbRn6GFP_Δ46-52_ with HindIII and MVa1269I. The upstream integration region of the resultant plasmid was then modified by extending the *VSG221* 3’ UTR sequence to include the native polypyrimidine tract as described for pbRn6.M1.6_WT_.

The cell lines Δ221^ES^121_WT_221^Tet^ and Δ221^ES^121_N46-48_221^Tet^ were generated by transfecting the single-expressor cell lines Δ221^ES^121_WT_ and Δ221^ES^121_N46-48_ with the overexpression plasmid pRS.221 (a modified pRS121(Batram *et al*., 2014) where *VSG121* was replaced with VSG221) after NotI linearisation.

The overexpression cell lines for EP1-eYFP overexpression (Δ221^ES^121_WT_EP1^Tet^ and Δ221^ES^121_N46-48_EP1^Tet^) were generated by transfecting the single-expressor cell lines Δ221^ES^121_WT_ and Δ221^ES^121_N46-48_ with the overexpression plasmid pRS.EP1::eYFP full3’ UTR. The plasmid was generated by a fusion PCR approach. Briefly, two PCR amplified fragments from the plasmid pGAPRONEΔLII_EP1::eYFP (Engstler and Boshart, 2004). Fragment 1 (EP1:eYFP with part of the EP1 3’ UTR, flanked by HindIII and the LII region) was amplified with the primer pair MBS59/MBS60. Fragment 2 (part of the EP1 3’ UTR flanked by XhoI and LII region) was amplified using the primer pair MBS61/MBS62. The two fragments were joined in a PCR using the primer pair MBS59/MBS62. The PCR product obtained (EP1::eYFP_full3’ UTR) was cloned into pJET1.2 to obtain the plasmid pJET1.2_EP1::eYFP_full3’ UTR. The EP1::eYFP_full3’ UTR fragment was then excised from pJET1.2_EP1::eYFP_full3’ UTR using HindIII and XhoI and ligated into the pLEW82v4 vector linearised with the same restriction enzymes. The resultant plasmid pRS.EP1::eYFP_full3’ UTR was linearised with NotI for transfection.

To generate the overexpression cell lines VSG121_WT_ and VSG121_N46-48_, *T. brucei* 13-90 cells were transfected with NotI linearised plasmids pRS.121 (Batram *et al*., 2014) and pRS.121_N46-48_, respectively. pRS.121_N46-48_ was generated using the fusion PCR approach. Two PCR amplified fragments from pbRn6M1.6TGA46-48ACT.nPPT were joined together. Fragment 1 (VSG121 ORF and part of the 3’ UTR) was amplified using primers MBS71/MBS75. Fragment 2 (part of the 3’ UTR) was amplified using primers MBS72/MBS76. The two fragments were fused together in a PCR using primers MBS71/MBS72. The PCR product obtained (M1.6N46-48) was cloned into pJET1.2 to obtain the plasmid pJET1.2_M1.6N46-48. The M1.6N46-48 fragment was then excised from pJET1.2_M1.6N46-48 using HindIII and XhoI and ligated into pLEW82v4 vector linearised with the same restriction enzymes.

For the overexpression cell lines VSG118_WT_ and VSG118_N46-48_, *T. brucei* 13-90 cells were transfected with NotI linearised plasmids pRS.118 (a modified pRS.121 plasmid where *VSG221* was replaced with *VSG118*) and pRS.118_N46-48_, respectively. To generate pRS.118_N46-48_, mutagenesis PCR by a fusion PCR approach was used to introduce the mutation TGA46-48ACT into the VSG118 3’ UTR using the plasmid pRS.118 as template. Fragments 1 and 2 were amplified using the primer pairs MBS85/MBS86 and MBS87/MBS88, respectively. The two fragments were then fused with primers MBS85/MBS88. The PCR product obtained (M1.5N46-48) was cloned in pJET1.2 to obtain the plasmid pJET1.2_M1.5N46-48. The M1.6N46-48 fragment was then excised from pJET1.2_M1.5N46-48 using HindIII and XhoI and ligated into the pLEW82v4 vector linearised with the same restriction enzymes.

### Differentiation of monomorphic bloodstream trypanosomes to the procyclic form

Differentiation of monomorphic bloodstream trypanosomes to the procyclic form was carried out *in vitro*. BSF trypanosomes were grown to a density of 1.5 x 10^6^ cells/ml in HMI-9 medium. 2.5 x 10^7^ cells were harvested by centrifugation (1,400x *g* for 10 min at room temperature (RT)) and resuspended in 5 ml of Differentiating Trypanosome Medium (DTM) containing 6 mM cis-aconitate (Overath, Czichos and Haas, 1986). The parasites were then cultivated at 27 °C and 5 % CO_2._ The cells were diluted with SDM-79 medium supplemented with 10 % heat-inactivated FCS when the cell density was above 2 x 10^7^ cells/ml.

### RNA isolation and mRNA quantification

Total RNA was extracted from 1 x 10^8^ trypanosome cells using the Qiagen RNeasy Mini Kit (Qiagen, Netherlands) following the manufacturer’s instructions. Poly(A)+ mRNA on the other hand was isolated from total RNA using the Oligotex mRNA Mini Kit (Qiagen, Netherlands) following the manufacturer’s protocol. *VSG* mRNA levels were quantified using RNA dot blots as described (Batram *et al*., 2014).

### N6-methyladenosine Immunoblotting

Immunoblotting to detect m^6^A modification was carried out as described by (Viegas *et al*., 2022) with slight modifications. DNase I-treated total RNA (2 μg) or poly(A)+ mRNA (50 ng) samples were mixed with a 5x RNA loading buffer (30.8 % (v/v) formamide, 2.6 % (v/v) formaldehyde, 20 % (v/v) glycerol, 0.2 % (w/v) bromophenol blue, 4 mM EDTA pH 8) in a ratio of 1 volume 5x loading buffer to 4 volumes of RNA sample. Samples were denatured by heating (70 °C for 5 min) and immediately transferred onto ice. The samples were then loaded and resolved on a 1.4 % denaturing agarose gel (1.4 % agarose, 6.3 % formaldehyde, 1x MOPS buffer) at 100 V until the samples had migrated halfway across the gel. The RNA was transferred overnight to a Hybond-N+ nylon membrane (GE Healthcare) by upward capillary transfer with 20x SSC buffer. The RNA was then UV-crosslinked to the membrane (Stratalinker 2400, autocrosslink: 120 mJ/cm^2^). Membranes were stained with 0.02 % methylene blue diluted in 0.3 M sodium acetate (pH 5.5) for 5 min and washed in RNase-free water. After imaging, the methylene blue was removed by incubation in destaining solution (0.2x SSC, 1 % SDS) and washed 3 times in PBST (PBS pH 7.4 with 0.2 % Tween-20). The membranes were then blocked by incubation in 5 % skimmed milk in PBST for 1 h followed by overnight incubation with 1 µg/ml mouse anti-m^6^A antibody (1:1000 in 2.5 % skimmed milk in PBST) at 4 °C. Membranes were washed three times in PBST and then incubated with HRP-conjugated anti-mouse IgG diluted 1:10000 in 2.5 % skimmed milk in PBST for 1 h at room temperature. Membranes were again washed three times in PBST (10 min each) and the signal developed using a Western Lightning Plus-ECL, Enhanced Chemiluminescence Substrate kit (PerkinElmer). Images were acquired using the iBright FL1000 imaging system (Thermo Fisher Scientific).

### Protein analysis

Protein amounts were quantified using either western blots or protein dot blots. For western blots protein samples from 1 x 10^6^ trypanosome cells were separated on 12.5 % sodium dodecyl sulphate (SDS)-polyacrylamide gels and transferred onto nitrocellulose membranes (GE Healthcare Life Sciences). For protein dot blots, 3 μl of protein samples containing 6 x 10^5^ cell equivalents were loaded onto nitrocellulose membranes fixed in a dot blotter. The membranes were blocked by incubation with 5 % milk powder in PBS for 1 h at room temperature (RT) or overnight at 4 °C. Primary antibodies (rabbit anti-VSG221; 1:5000, rabbit anti-VSG121; 1:2000, rabbit anti-VSG118; 1:10000 and mouse anti-PFR antibody (L13D6, 1:20) (Kohl, Sherwin and Gull, 1999)) were then applied in PBS/1 % milk/0.1 % Tween-20 solution for 1 h at RT. After four washes (5 min each) with PBS/0.2 % Tween-20, IRDye 800CW-conjugated goat-anti-rabbit and IRDye 680LT-conjugated goat-anti-mouse secondary antibodies (1:10000; LI-COR Biosciences) were applied in PBS/1 % milk/0.1 % Tween-20 solution for 1 h at RT in the dark. The membranes were washed four times (5 min each in the dark) with PBS/0.2 % Tween-20 followed by a final 5 min wash with PBS. Blots were analysed using a LI-COR Odyssey system (LI-COR Biosciences).

## Acknowledgement

We thank Prof. Susanne Kramer for technical advice and discussions on the manuscript. We are grateful to Prof. Mark Carrington for discussions on the project during MB-S’s doctoral studies. MB-S was supported by a grant from the German Excellence Initiative awarded to the Graduate School of Life Sciences, University of Würzburg. ME was supported by DFG grants EN305, SPP1726, and GRK2157, GIF grant I-473-416.13, the EU ITN Physics of Motility, and the BMBF NUM Organo-Strat. ME is a member of the Wilhelm Conrad Röntgen Center for Complex Material Systems (RCCM).

## Author contributions

M.B.-S., C.B., H.Z., N.G.J., and M.E. conceived and designed the experiments; M.B.-S., C.B., and H.Z. conducted the experiments; M.B.-S., C.B., N.G.J., and M.E. analysed and interpreted the results; M.B.-S., N.G.J., and M.E. wrote the initial manuscript. All the authors reviewed the manuscript.

## Notes

### Competing Interest Statement

The authors have declared no competing interest.

### Summary of Updates

The revision includes minor changes in nomenclature, new citations and few new points in the discussion.

