## Supplemental Material for "The dual role of the 16mer motif within the 3’ untranslated region of the variant surface glycoprotein of *Trypanosoma brucei*"

### Supplementary data

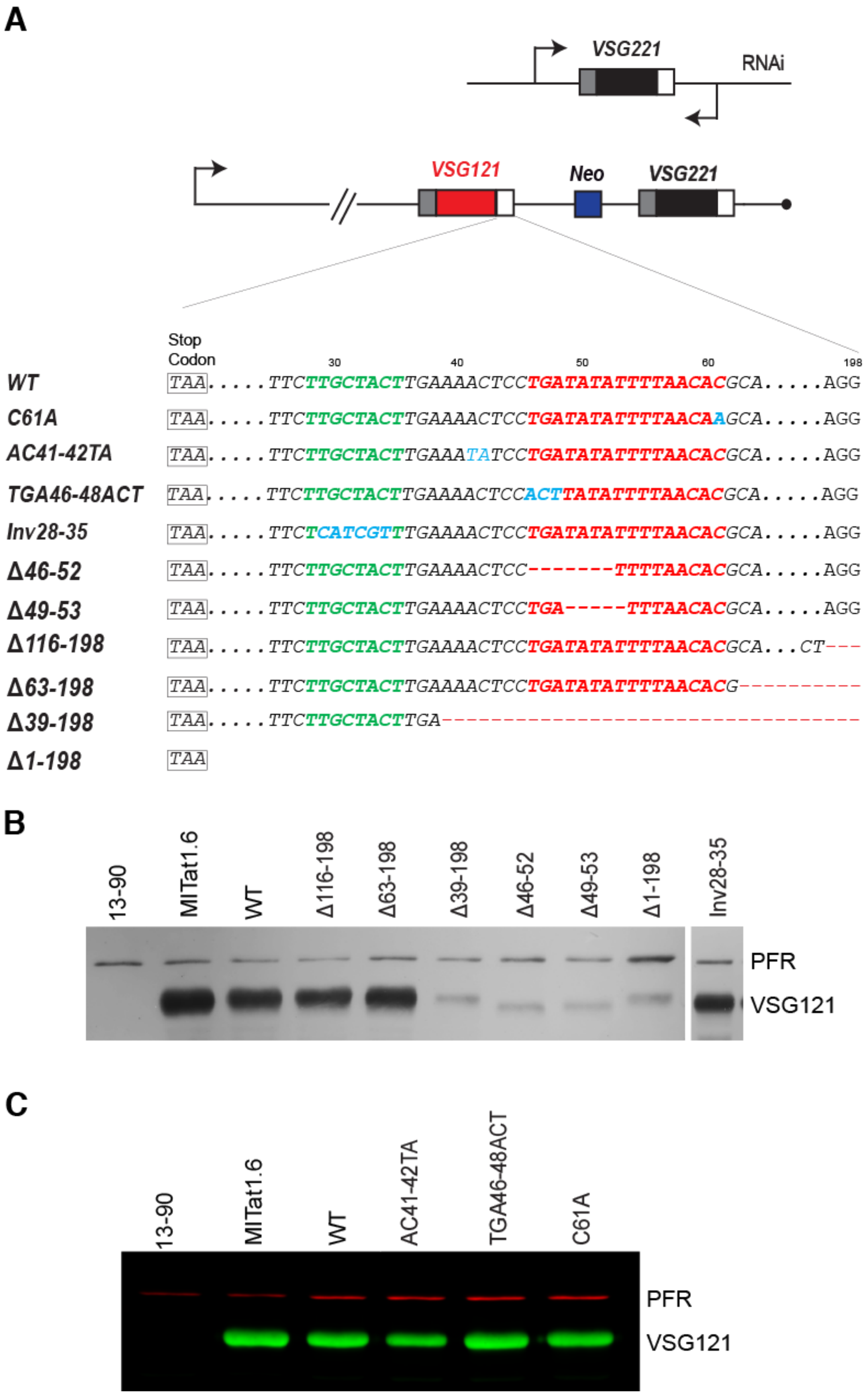


**Figure S1. Mutational analysis of the VSG121 3´ UTR.** (A) Schematic of the inducible VSG221 RNAi double-expressor cell line with specific mutations of the VSG121 3´ UTR. NEO: Neomycin resistance cassette. (B, C) Western blots showing VSG121 protein levels upon mutations in the VSG121 3´ UTR. Note that the 3'UTR mutants Δ46-52 and Δ49-53 were tested using an N-glycosylation deletion mutant of VSG 121 (Hartel et al., 2016). This explains the slightly reduced size of the protein.

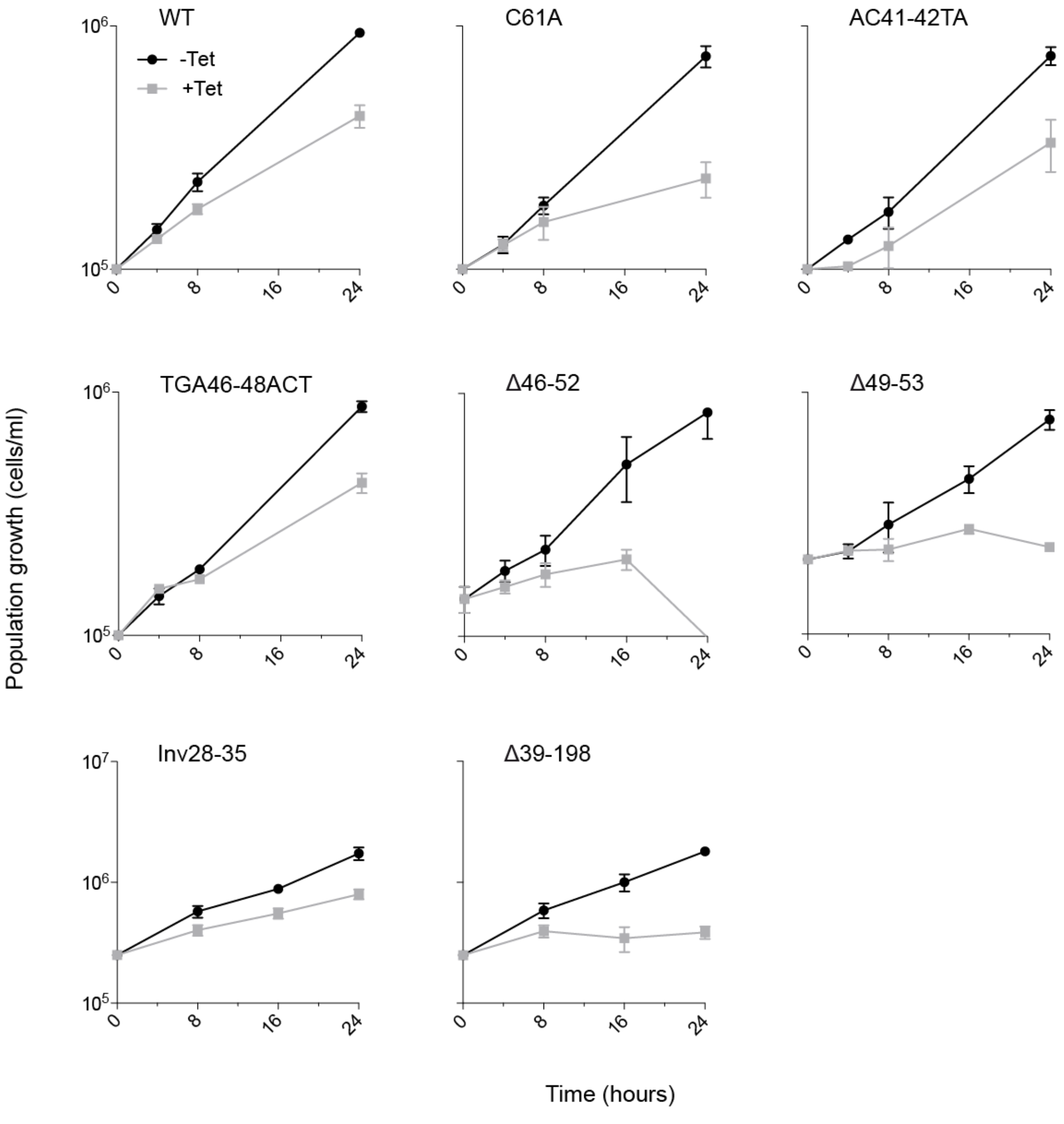


**Figure S2. VSGs with tolerated mutations support cell growth upon RNAi-mediated depletion of VSG221.** Cumulative growth curves of double-expressor cells upon RNAi-mediated depletion of VSG221. Mutations Δ46-52, Δ49-53 and Δ39-198 are not tolerated. They do not support growth as can be seen in the cell density at 24h compared to the Wt and other mutants.

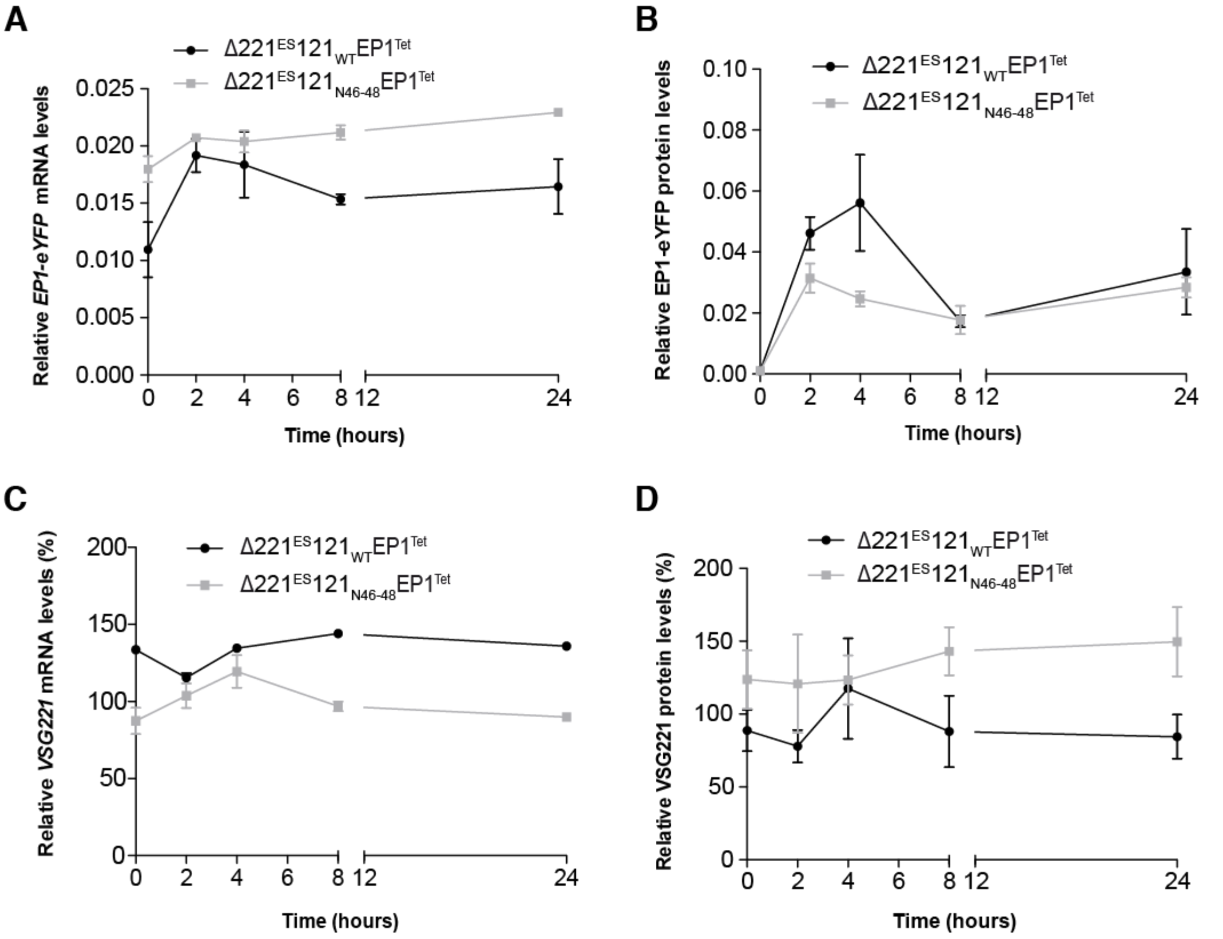


**Figure S3. Overexpression of EP1-eYFP in $\Delta221^{ES}121_{WT}$ and $\Delta221^{ES}121_{N46-48}$ single-expresser cell lines**. *VSG221* mRNA (A) and VSG221 protein (C) monitored in $\Delta221^{ES}121_{WT}EP1^{Tet}$ and $\Delta221^{ES}121_{N46-48}EP1^{Tet}$ cells during the course of EP1-eYFP overexpression. *VSG* mRNA and VSG protein levels were quantified from RNA and protein dot blots, respectively. *VSG* mRNA was normalised to *tubulin* mRNA and the protein amounts normalised to PFR. The VSG expression levels are given relative to levels in the parental 13-90 (VSG221) cells and expressed as mean ± standard error of the mean (SEM) of three clonal cell lines, respectively. *EP1-eYFP* mRNA (B) and EP1-eYFP protein (D) monitored in $\Delta221^{ES}121_{WT}EP1^{Tet}$ and $\Delta221^{ES}121_{N46-48}EP1^{Tet}$ cells during the course of EP1-eYFP overexpression. *EP1-eYFP* mRNA and EP1-eYFP protein were normalised to *tubulin* mRNA and PFR, respectively. Values are expressed as mean ± standard error of the mean (SEM) of three clonal cells each.

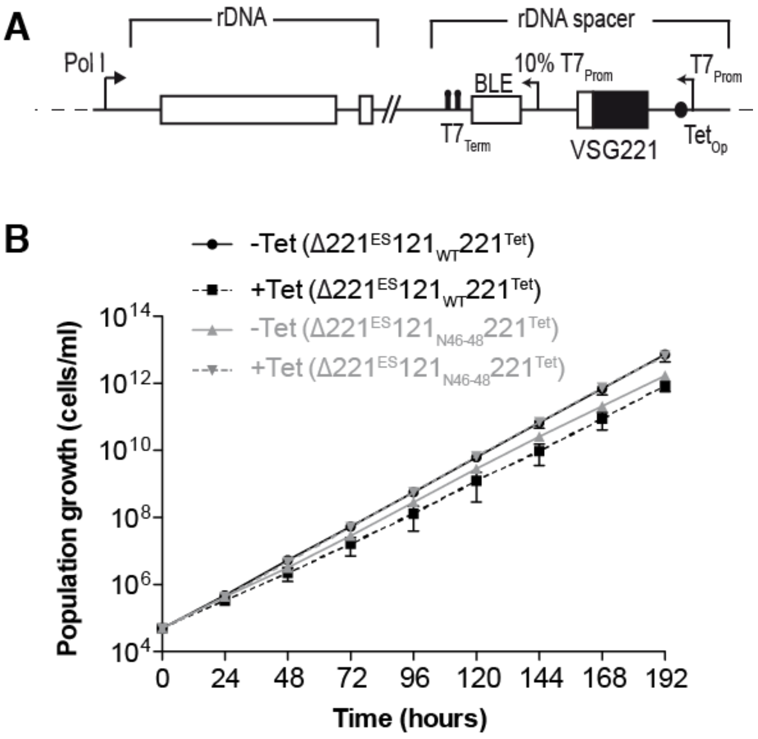


**Figure S4. Overexpression of VSG221 in Δ221$^{ES}$121$_{WT}$ and Δ221$^{ES}$121$_{N46-48}$ single-expresser cell lines.** (A) Schematic of the ectopic overexpression system (adapted from Batram et al., 2014). Cumulative growth curves of Δ221$^{ES}$121$_{WT}$221$^{Tet}$ (black) and Δ221$^{ES}$121$_{N46-48}$221$^{Tet}$ (grey) cell lines over the course of VSG221 overexpression.
